# Quantifying genetic heterogeneity between continental populations for human height and body mass index

**DOI:** 10.1101/839373

**Authors:** Jing Guo, Andrew Bakshi, Ying Wang, Longda Jiang, Loic Yengo, Michael E Goddard, Peter M Visscher, Jian Yang

**Affiliations:** Institute for Molecular Bioscience, The University of Queensland, Brisbane, Queensland 4072, Australia; Monash Partners Comprehensive Cancer Consortium, Monash Biomedicine Discovery Institute Cancer Program, Prostate Cancer Research Group, Department of Anatomy and Developmental Biology, Monash University, Clayton, Victoria 3800, Australia; Queensland Brain Institute, The University of Queensland, Brisbane, Queensland 4072, Australia; Faculty of Veterinary and Agricultural Science, University of Melbourne, Parkville, Victoria, Australia; Biosciences Research Division, Department of Economic Development, Jobs, Transport and Resources, Bundoora, Victoria, Australia; Institute for Advanced Research, Wenzhou Medical University, Wenzhou, Zhejiang 325027, China

## Abstract

Genome-wide association studies (GWAS) in samples of European ancestry have identified thousands of genetic variants associated with complex traits in humans. However, it remains largely unclear whether these associations can be used in non-European populations. Here, we seek to quantify the proportion of genetic variation for a complex trait shared between continental populations. We estimated the between-population correlation of genetic effects at all SNPs (*r*_*g*_) or genome-wide significant SNPs (*r*_*g(GWS)*_) for height and body mass index (BMI) in samples of European (EUR; *n* = 49,839) and African (AFR; *n* = 17,426) ancestry. The 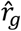 between EUR and AFR was 0.75 (s. e. = 0.035) for height and 0.68 (s. e. = 0.062) for BMI, and the corresponding 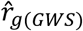 was 0.82 (s. e. = 0.030) for height and 0.87 (s. e. = 0.064) for BMI, suggesting that a large proportion of GWAS findings discovered in Europeans are likely applicable to non-Europeans for height and BMI. There was no evidence that 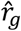 differs in SNP groups with different levels of between-population difference in allele frequency or linkage disequilibrium, which, however, can be due to the lack of power.

## Introduction

Most traits and common diseases in humans are complex because they are influenced by many genetic variants as well as environmental factors^1,2^. Genome-wide association studies (GWASs) have discovered >70,000 genetic variants associated with human complex traits and diseases^3,4^. However, most GWASs have been heavily biased toward samples of European (EUR) ancestry (~79% of the GWAS participants are of EUR descent)^5^. Progress has been made in recent years in uncovering the genetic architecture of traits and diseases in a broader range of populations^6–11^. Given the population genetic differentiation among worldwide populations^5,12–15^, the extent to which the associations discovered in EUR populations can be used in non-EUR such as Africans (AFR) and Asians remains unclear. Genetic correlation (*r*_*g*_) is the correlation between the additive genetic values of two traits in a population^16^. However, by definition, we cannot observe the trait in AFR and EUR in the same individuals. Therefore, *r*_*g*_ is better defined by the correlation between the additive effects of causal variants in the two populations. *r*_*g*_ can be less than 1 due to genotype by environment interactions if the two populations are in different environments. Unfortunately, not all the causal variants for complex traits are known so we estimate *r*_*g*_ based on the correlation between the apparent effects of genetic markers such as SNPs. This can be estimated by using the genomic relationship matrix (GRM) among all the individuals or, if only summary data is available, the correlation between estimated SNP effects^13,17–19^. *r*_*g*_ estimated from SNPs can be less than that based on causal variants if the LD between causal variants and SNPs differs between the populations. Galinsky *et al.*^14^ estimated this effect using simulation and found it to be small but this conclusion may not apply to rare causal variants.

Previous trans-ethnic genetic studies have shown that the estimates of *r*_*g*_ at common SNPs (e.g., those with minor allele frequencies (MAF) > 0.01) between EUR and East Asian (EAS) populations are high for inflammatory bowel diseases (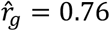 with a standard error (s.e.) of 0.04 for Crohn’s disease and 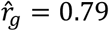 with s. e. = 0.04 for ulcerative colitis)^20^ and bipolar disorder 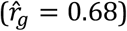^21^ and modest for rheumatoid arthritis (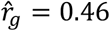 with s. e. = 0.06)^13^ and major depressive disorder (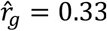 with a 95% confidence interval (CI) of 0.27-0.39)^22^. If the between-population *r*_*g*_ for a trait estimated from SNPs is not unity, then it is of interest to know whether the between-population genetic heterogeneity differs at SNPs with different levels of between-population difference in allele frequency (i.e., Wright’s fixation index^23^, *F*_ST_) or LD, and whether the between-population *r*_*g*_ estimated from all common SNPs (MAF > 0.01) can be used to measure the correlation of genetic effects between populations at the genome-wide significant SNPs. Answers to these questions are important to inform the design of gene mapping experiments^24–28^, the genetic risk prediction of complex diseases^5,29^ in the future in non-EUR populations and the detection of signatures of natural selection that has resulted in genetic differentiation among worldwide populations. In this study, we focus on estimating the correlation of genetic effects at all SNPs (denoted by *r*_*g*_) between continental populations using a bivariate GREML analysis^30^ (treating the phenotypes in the two populations as different traits) for two model complex traits, i.e., height and body mass index (BMI). We investigate the influence of the between-population differences in allele frequencies or LD on the between-population genetic heterogeneity. To do this, we first used genome-wide SNP genotype data to estimate *r*_*g*_ between AFR and EUR populations for height and BMI. We also estimated the correlation of genetic effects between continental populations at the genome-wide significant SNPs (*r*_*g(GWS)*_) identified from an EUR GWAS using the bivariate GREML method^30^ or a summary level data-based method^31^. We then examined whether the between-population genetic overlap is enriched (or depleted) at the SNPs with stronger between-population differentiation in allele frequency or LD.

## Results

### Genetic correlation (*r*_*g*_) between worldwide populations for height and BMI

We used GWAS data on 49,839 individuals of EUR ancestry from the UK Biobank (UKB) and 17,426 individuals of AFR ancestry from multiple publicly available datasets including the UKB (Supplementary Fig. 1; Methods). Note that we used only ~50K EUR individuals from the UKB for the ease of computation. All the individuals were not related in a sense that the estimated pairwise genetic relatedness was < 0.05 within a population. The EUR genotype data were imputed by the UKB (version 3) using the Haplotype Reference Consortium (HRC) and UK10K imputation reference panel^32^. We imputed the AFR data to the 1000 Genomes Project (1000G) reference panel (Methods). After quality control (QC), 1,018,256 HapMap3 SNPs with MAF >0.01 in both the two data sets were retained for analysis (Methods). We first used the bivariate GREML approach^30^ to estimate *r*_*g*_ between populations as well as the SNP-based heritability 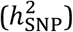 in each population for height and BMI. It has been shown in Galinsky *et al.*^14^ that the estimate of *r*_*g*_ from a between-population bivariate GREML analysis is equivalent to the correlation of genetic effect at all SNPs. The GRM used in our bivariate GREML analysis was computed using two different strategies: 1) SNP genotypes standardized using allele frequencies estimated from a combined sample of the two populations (denoted as GRM-average); 2) SNP genotypes standardized using allele frequencies estimated from each population specifically (denoted as GRM-specific; Methods). The 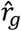 based on GRM-specific was 0.75 (s. e. = 0.035) for height and 0.68 (s. e. = 0.062) for BMI, suggesting strong genetic overlap between EUR and AFR for both height and BMI (Table 1). The 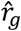 between EUR and AFR for height was very similar to that between EUR and SAS estimated from the UKB data reported in Galinsky *et al.* (0.77 with s. e. = 0.26)^14^. We did not observe a substantial difference in 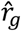 between the analyses based on GRM-average (Supplementary Table 1) and GRM-specific (Table 1). The 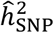 in EUR and AFR from the bivariate GREML analysis were 0.50 (s. e. = 0.0077) and 0.39 (s. e. = 0.024) for height, and 0.25 (s. e. = 0.0080) and 0.22 (s. e. = 0.025) for BMI, respectively (Table 1), highly consistent with those from the univariate GREML analysis^33^ where the corresponding estimates were 0.50 (s. e. = 0.0078) and 0.40 (s. e. = 0.026) for height, and 0.25 (s. e. = 0.0080) and 0.23 (s. e. = 0.025) for BMI. It is of note that the height 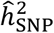 in EUR was significantly larger than that in AFR (*P* = 1.3 × 10^−4^), which is consistent with the result from a recent study in European-Americans and African-Americans^15^, presumably because the causal variants in non-Europeans, especially those with MAF <0.01, were less well tagged by the SNPs on the SNP arrays compared to those in Europeans. Such a difference was much smaller and not statistically significant for BMI (*P* = 0.35), which can be partly explained by that the imperfect tagging is proportional to trait heritability^34^. We further estimated *r*_*g*_ between EUR and EAS for BMI by a summary-data-based *r*_*g*_ approach^13^ using summary statistics from the GIANT consortium (*n* = 253,288)^35^ and the Biobank Japan project (BBJ, *n* = 158,284)^10^ (note that the GWAS data with comparable sample size for EAS and the BBJ summary-level data for height were not available to us). The 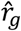 between EUR and EAS was 0.80 (s. e. = 0.037) for BMI, which was also significantly different from 1 (*P* = 8.36 × 10^−8^), in line with the estimate (0.75, s. e. = 0.023) from Martin *et al.*^5^ based on GWAS summary data from the UKB and BBJ.

**Table 1.**
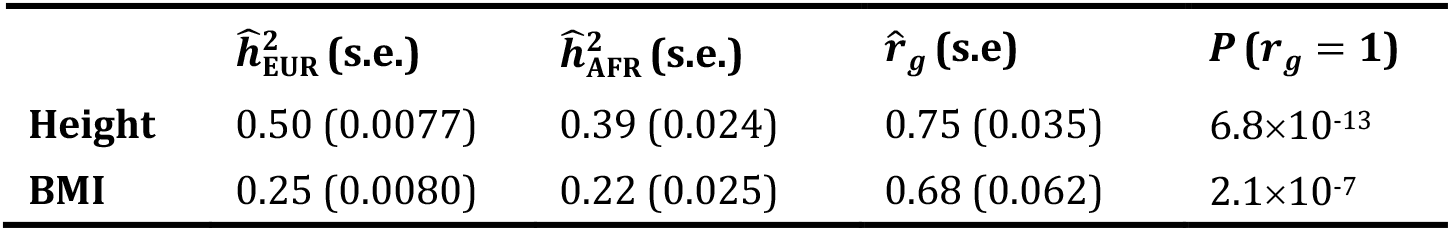
Estimated 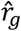 between EUR and AFR using HapMap3 SNPs based on GRMs-specific for height and BMI.

### Correlation of SNP effects between populations at the top associated SNPs

We have quantified above the between-population *r*_*g*_ for height and BMI using all HapMap3 SNPs with MAF >0.01. The estimates were high but statistically significantly smaller than 1 (Table 1), suggesting there is a between-population genetic heterogeneity for both traits. We know from a previous study that 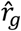 estimated from all SNPs is close to the estimated causal effect correlation 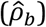 between EUR and SAS^14^. We then sought to ask whether the estimated *r*_*g*_ from all SNPs is consistent with that estimated at genome-wide significant SNPs identified in EUR (i.e., *r*_*g(GWS)*_). We estimated *r*_*g(GWS)*_ between EUR and AFR using the recently developed method^31^ that can estimate SNP effect correlation using summary data accounting for errors in the estimated SNP effects (Methods). We used the trait-associated SNPs identified in previous GWAS meta-analyses conducted by the GIANT consortium^35,36^ (with SNP effects re-estimated in our AFR and EUR samples to avoid biases due to the winner’s curse; see Methods). There were 538 and 57 nearly independent SNPs for height and BMI respectively at *P* < 5.0 × 10^−8^ selected from clumping analyses (LD *r*^2^ threshold = 0.01 and window size = 1Mb) of the GIANT summary data (Methods)^37^. To avoid potential bias in estimating *r*_*g(GWS)*_ due to remaining LD among these sentinel SNPs, we did an additional round of clumping using a window size of 10Mb (Methods) and obtained 531 and 56 SNPs for height and BMI respectively. We call these the sentinel SNPs hereafter.

We first estimated *r*_*g(GWS)*_ between our EUR sample and GIANT as a “negative control”; the estimate was 0.98 (s. e. = 0.0045) for height and 0.99 (s. e. = 0.0069) for BMI, suggesting no significant differences in SNP effects between the GIANT (a meta-analysis of samples of EUR ancestry) and our sample of EUR participants from the UKB (Figure 1). We then estimated *r*_*g(GWS)*_ between EUR and AFR (SNP effects re-estimated in our samples). We found an estimate of 0.81 (s. e. = 0.032) for height (Figure 1a) and of 0.94 (s. e. = 0.049) for BMI (Figure 1b). Since individual-level data were available in our EUR and AFR samples, we performed a bivariate GREML analysis to estimate *r*_*g(GWS)*_ only using the sentinel SNPs (Methods); the estimate was 0.82 (s. e. = 0.030) for height and 0.87 (s. e. = 0.064) for BMI, similar to the corresponding estimates using the summary data above. Moreover, summary data-based 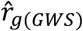 between EUR (SNP effects re-estimated in this study) and EAS (SNP effects from the BBJ data^38^) was 0.90 (s. e. = 0.043) for BMI. All these results suggest that a large proportion of GWAS findings discovered in Europeans are likely replicable in non-Europeans for the two traits (see below for more discussion). In addition, 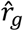 estimated using all SNPs was largely consistent with 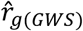 for height, but some differences have been observed for BMI (see below for discussion).

**Figure 1.**
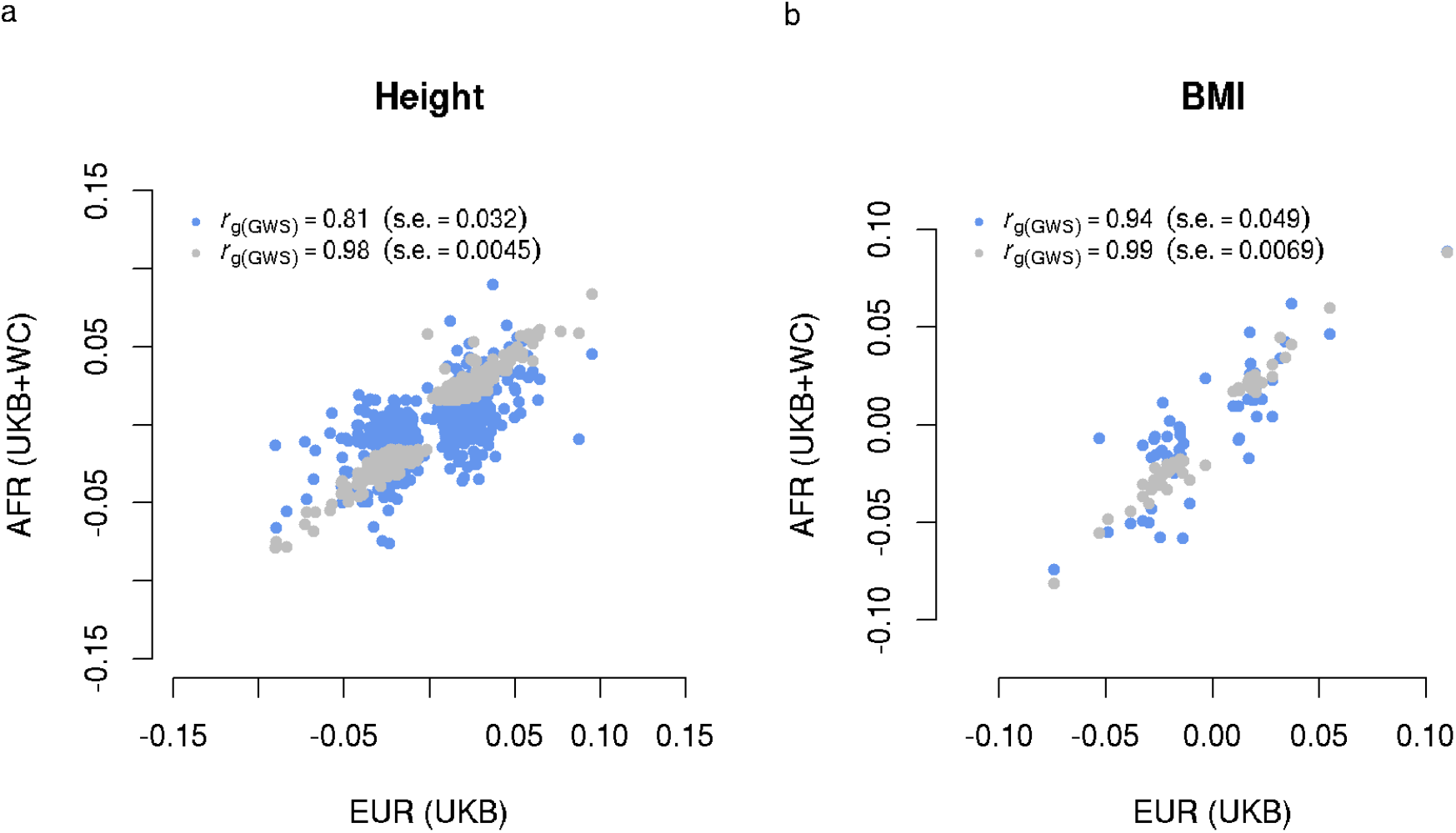
Estimated genetic effect correlation between AFR and EUR for height (a) and BMI (b) at genome-wide significant SNPs. The near-independent trait-associated SNPs were discovered in GIANT with their effects re-estimated in our EUR (*n* = 456,422) and AFR (*n* = 23,355) data. The blue dots show a comparison of SNP effects between EUR and AFR and the grey ones show the comparison within EUR (i.e., GIANT vs. EUR-UKB).

### Genetic correlation estimated at SNPs stratified by population difference in allele frequency or LD

If there is an effect of the between-population differences in allele frequencies on the between-population genetic heterogeneity for a trait, we hypothesised that the estimate of *r*_*g*_ at SNPs with higher *F*_ST_ is different from that at SNPs with lower *F*_ST_. To test this, we first calculated the *F*_*ST*_ values of the HapMap3 SNPs between EUR and AFR. To avoid difference in within-population allele frequency or LD between the two *F*_ST_ groups, we divided the SNPs into a large number of bins according to their allele frequencies and LD scores in each population and then stratified the SNPs into two groups with equal number by their *F*_*ST*_ values in each MAF-LD bin (Methods). We show that there was no difference in allele frequency or LD score between the two *F*_*ST*_ groups after applying this SNP-binning strategy (Supplementary Fig. 2). We performed a two-component bivariate GREML analysis (based on GRM-specific) to estimate *r*_*g*_ in each *F*_*ST*_ group and found no significant difference in 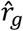 between the two *F*_*ST*_ groups for both traits although the standard errors of 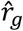 were large (Table 2). Even if our previous study has shown that height increasing alleles are more frequent in EUR than AFR^39^, which might explain the mean difference in height phenotype between EUR and AFR, the result reported here suggests that the population differentiation of frequencies of the height-associated SNPs does not seem to affect the genetic correlation between populations. Nevertheless, it is possible that there is a difference in *r*_*g*_ between the two *F*_*ST*_ groups but the power of this study is not large enough to detect it.

**Table 2.** Difference of the estimated 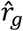 for EUR-AFR between SNP sets stratified by allele frequency- and LD-matched *F*_*ST*_ (and LDCV) for height and BMI respectively.

| | $F_{ST}$ stratified | | | LDCV stratified | | |
| --- | --- | --- | --- | --- | --- | --- |
| | $\hat{r}_{g-l}$ (s.e.) | $\hat{r}_{g-h}$ (s.e.) | $P_{\text{difference}}$ | $\hat{r}_{g-l}$ (s.e.) | $\hat{r}_{g-h}$ (s.e.) | $P_{\text{difference}}$ |
| **Height** | 0.84 (0.15) | 0.68 (0.099) | 0.509 | 0.92 (0.12) | 0.59 (0.087) | 0.076 |
| **BMI** | 0.73 (0.19) | 0.62 (0.23) | 0.785 | 0.82 (0.29) | 0.63 (0.15) | 0.640 |

l and h indicate the SNP group with lower $F_{ST}$ (or LDCV) and higher $F_{ST}$ (or LDCV), respectively.

We applied the same SNP-binning strategy to test whether the estimate of genetic correlation differs when the SNPs are ascertained by difference in LD between populations (Supplementary Fig. 3). We used a metric called LDCV (i.e., coefficient of variation of the LD scores across populations) proposed in a previous study^39^ to measure the differentiation of LD-score between EUR and AFR for each SNP (Methods). We stratified the SNPs into two LDCV groups with no difference in MAF or LD score between the groups in each individual population using the approach described above (Methods; Supplementary Fig. 4) and estimated *r*_*g*_ by a two-component bivariate GREML analysis. We found no significant difference in the estimate of 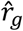 between the two LDCV groups (Table 2), which does not support a significant role of LD difference in the between-population genetic heterogeneity at common SNPs but also could be due to the lack of power if the difference in *r*_*g*_ between the two LDCV groups is very small.

## Discussion

In this study we showed a substantial genetic overlap at HapMap3 SNPs (MAF > 0.01) for height and BMI between EUR and AFR (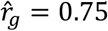 with s. e. = 0.035 for height and 0.68 with s. e. = 0.062 for BMI; Table 1) from a cross-population bivariate GREML analysis of individual-level genotype data^30^ and between EUR and EAS (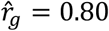 with s. e. = 0.037 for BMI) by a summary-data-based approach^13^. All these estimates were significantly smaller than 1 (Table 1), suggesting some genetic heterogeneity between populations for both traits. We then used the recently developed *r*_*b*_ approach^31^ that is able to estimate the correlation of SNP effects between populations accounting for estimation errors in estimated SNP effects (Figure 1), and confirmed the estimates by a bivariate GREML analysis using individual-level data. The bivariate GREML estimate of *r*_*g*_ at the sentinel SNPs between EUR and AFR was marginally larger than the estimate of *r*_*g*_ for height (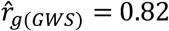 with s. e. = 0.030 vs. 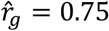 with s. e. = 0.035; *P* = 0.13), but the difference was larger for BMI (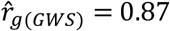 with s. e. = 0.064 vs. 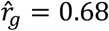 with s. e. = 0.062; *P* = 0.032), which may due to a difference in genetic architecture between the two traits and/or the relatively small number of sentinel SNPs used for BMI. The estimated strong correlation in SNP effect between populations is in line with the finding from previous studies that GWAS results from EUR population are largely consistent with those from non-EUR populations for a certain number of complex traits^17,40–45^. However, the extent to which the EUR-based GWAS findings can be replicated in non-EUR populations can be trait-dependent, given the estimates of genetic correlation varied across different traits^5,22^. We also attempted to quantify the effect of population differentiation in SNP allele frequencies on the between-population genetic heterogeneity by comparing 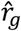 estimated from SNPs with higher *F*_*ST*_ to that estimated from SNPs with lower *F*_*ST*_ but found no significant difference in 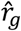 between the two *F*_*ST*_ groups (Table 2). In addition, it should be noted that differences in SNP effects between populations could reflect the differences in causal effects and/or LD between SNPs and causal variants. Our estimated genetic effect correlation at all SNPs between EUR and AFR for height (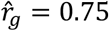 with s. e. = 0.035; Table 1) was largely consistent with the causal effect correlation between EUR and SAS (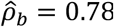, s. e. = 0.26) estimated in a previous study^14^. Although the standard error of 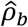 is large, the causal effect correlation between EUR and AFR is similar to that between EUR and SAS. Then, the results seem to imply that, on average, the extent to which the difference in SNP effects between populations due to the difference in LD is unlikely to be large for common SNPs. This implication is consistent with our LDCV partitioning analysis which showed no significant difference in 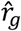 between common SNPs with higher and lower LDCV (Table 2). However, it should be noted that LDCV may differ from the between-population difference in LD between SNPs and causal variants.

In summary, our study confirmed a large estimate of genetic correlation at common SNPs between worldwide populations for height^14^ and showed a similar level of between-population genetic correlation for BMI. We observed that the estimate of SNP effect correlation at the genome-wide significant SNPs was only marginally larger than the estimate of genetic correlation using all SNPs for height but the difference was more pronounced for BMI. We caution that the difference between 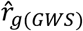 and 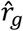 needs to be quantified in higher precision and the extent to which the between-population genetic heterogeneity for a trait due to differences in allele frequency and LD need to be tested in data sets with larger sample sizes in the future. Moreover, an observed between-population genetic heterogeneity for a complex trait could also be due to the interactions between genetic (G) and environmental (E) factors. The genotype-by-environment interaction component would be partially eliminated in *r*_*g*_ estimation in the study design where two populations differ in genetic ancestry but live in the same environment conditions. We acknowledge that all the conclusions are restricted to common SNPs. The between-population genetic heterogeneity for complex traits at rare variants (or the variants that are rare in one population but common in another) remains to be explored with whole-genome sequence data in large samples^46^. Nevertheless, all our results are consistent with the conclusion that most GWAS findings at common SNPs from EUR populations are largely applicable to non-EUR for height and BMI for variant/gene discovery purposes. However, cautions are required for phenotype (or disease risk) prediction given the limited accuracy of genetic prediction using EUR-based GWAS results in non-EUR populations as demonstrated in recent studies^5,29^.

## Methods

### Data

GWAS data of 456,422 individuals of European ancestry were from the UKB (EUR-UKB). GWAS data of 24,077 individuals of African ancestry were from the UKB (AFR-UKB, *n* = 8,230), the Women’s Health Initiative (WHI; *n* = 7,480), and the National Heart, Lung, and Blood Institute’s Candidate Gene Association Resource (CARe) including ARIC, JHS, CARDIA, CFS and MESA (*n* = 8,367)^47^. QC of the UKB SNP genotypes had been conducted by the UKB QC team^32^ and the EUR-UKB data had been imputed to the HRC and UK10K reference panel. For the EUR-UKB imputed data (hard-call genotypes), we filtered out SNPs with missing genotype rate >0.05, MAF <0.01, imputation INFO score <0.03 or *P*-value for HWE test <10^−6^. We cleaned the WHI and CARe (AFR-WC) genotype data following the protocol provided by the dbGaP data submitters. We further removed SNPs with SNP call rate <0.95, MAF <0.01 or Hardy-Weinberg Equilibrium (HWE) test *P* <0.001, and removed individuals with sample call rate <0.9. We imputed the AFR-UKB and AFR-WC data to the 1000G using IMPUTE2^48^, and applied the same filtering thresholds as above to the imputed data. We then combined the cleaned AFR-UKB and AFR-WC as one AFR data set. Since the AFR samples are ancestrally more heterogeneous than the EUR-UKB sample, we removed the AFR individuals whose PC1 or PC2 were more than 6 s.d. away from the mean of the AFR in 1000G in AFR-WC and AFR-UKB separately (the PC-based QC of the EUR-UKB sample was described in a previous study^49^). Only the SNPs in common with those in HapMap3 SNPs (*m* = ~1,018,000) were retained for analysis. We used GCTA^50^ to construct the GRM in each population based on all the HapMap3 SNPs and removed one of each pair of individuals with estimated genetic relatedness >0.05 in each population (retained 348,501 and 17,693 unrelated individuals in the EUR-UKB and AFR, respectively). These unrelated AFR individuals were a subset of the AFR samples after PC-based QC. The first 20 principal components (PCs) were derived from the GRM in each population. Phenotypes in each population were adjusted for covariates (i.e., age in AFR-WC, and age and assessment centre in EUR-UKB and AFR-UKB) in each gender group of each cohort and inverse-normal transformed after removing outliers that were 5 s.d. from the mean for height and 7 s.d. from the mean for BMI.

### GREML analyses to estimate 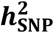 and *r*_*g*_ using all HapMap3 SNPs

To estimate 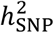 and cross-population *r*_*g*_ for each trait, we conducted a bivariate GREML analysis using all HapMap3 SNPs in the unrelated individuals (genetic relatedness <0.05). For the ease of computation, only 50,000 EUR individuals randomly sampled from the EUR-UKB data were included in the GREML analysis (all the AFR unrelated individuals were included in the analysis). To build the GRM for the bivariate GRM analysis (denoted by GRM-specific), the SNP genotypes were standardized based on the allele frequencies in a specific population (i.e., 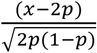 with *x* being coded as 0, 1 or 2 and *p* being the allele frequency in EUR, for example) using GCTA (--sub-popu option)^50^. The bivariate GREML analyses were then performed for height and BMI using the GRM-specific in a combined sample of EUR and AFR. The first 20 PCs generated from the GRM-specific were fitted as covariates in the bivariate GREML to control for population stratification. Only the samples that have both the genotype and phenotype data were included in the bivariate GREML analysis (*n* = 49,839 for EUR and *n* = 17,426 for AFR). We also performed the bivariate GREML analyses based on GRMs (and PCs thereof) for which the SNP genotypes were standardized using the allele frequencies computed from the combined sample of EUR and AFR. The bivariate GREML analyses were also performed to estimate *r*_*g(GWS)*_ using the GRM-specific built from the sentinel SNPs for both traits.

To compare the difference in 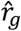 between SNP groups with higher and lower *F*_*ST*_, we computed *F*_*ST*_ between EUR and AFR for each SNP in GCTA (--fst option)^50^. We first split the SNPs into 125 bins according to their MAF in EUR and 125 bins based on the frequencies of the same alleles in AFR (125*125 frequency bins in total). We next split each frequency bin into 4 LD bins according to LD scores of the SNPs^34^ in EUR and 4 bins based on LD scores in AFR. We thereby obtained 250,000 (125*125*4*4) bins in total. We then equally divided the SNPs in each bin (*m* = 4 in most bins) into two groups according to the sorted *F*_*ST*_ values. There were a small number of bins with only 3 SNPs. For those bins, we randomly allocated 1 or 2 SNPs to the high-*F*_*ST*_ group and the remaining SNPs to the low-*F*_*ST*_ group. Finally, we combined the SNPs across all the bins with high and low *F*_*ST*_ respectively and computed the GRM-specific for each of the two SNP groups, and fitted the two GRMs jointly in a bivariate GREML analysis to estimate the between-population *r*_*g*_ and the population-specific 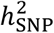 in each *F*_*ST*_ group for height and BMI. The first 20 PCs generated from the GRM-specific were fitted as covariates in the GREML analysis. The same strategy was applied to the LDCV stratification based on 250,000 bins including 20*20 frequency bins and 25*25 LD bins. The method to compute LDCV has been described elsewhere^39^.

### Testing the difference in 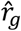 between SNP sets

We tested the difference in 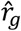 between two SNP sets (e.g., the two *F*_ST_-stratified SNP sets described above). We computed the *P*-value for the difference using a *χ*^2^ statistic with one degree of freedom, where 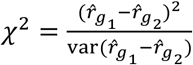 with 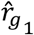 and 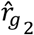 representing the estimates of the two SNP sets respectively, and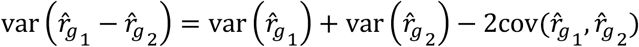. In the bivariate GREML analysis, *r*_*g*_ is defined as 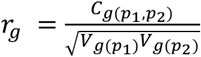 where 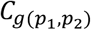 is genetic covariance between populations; 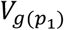 (or 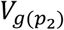) is the genetic variance in a population. The sampling variance of the estimate of *r*_*g*_ in a SNP set is

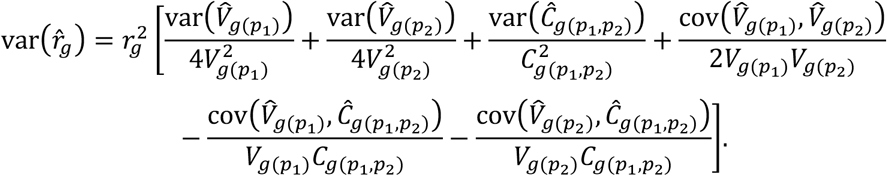

The sampling covariance of the estimates of *r*_*g*_ between two SNP sets is

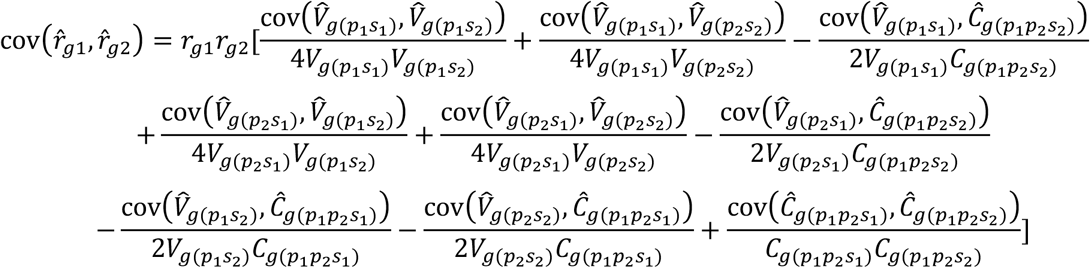

where the subscripts *s*_1_ (or *s*_2_) represents a SNP set. In practice, the parameters in the equations above can be replaced by their estimates to compute the estimates of 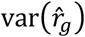 and 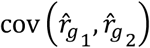.

### Estimation of SNP effect correlation between populations from GWAS summary data

We obtained the trait-associated SNPs for height and BMI from the GIANT meta-analyses^35,36^. We used the *r*_b_ method developed by Qi *et al.*^31^ to estimate the correlation of SNP effects between populations at the top associated SNPs accounting for sampling errors in the estimated SNP effects. To avoid bias due to ‘winner’s curse’, we re-estimated the SNP effects in our samples (independent from the samples used in the GIANT meta-analysis) using fastGWA^51^. Since fastGWA controls for relatedness^51^, we used all the samples passed QC (including close relatives) for the GWAS analysis (*n* = 456,422 for EUR and 23,355 for AFR after PC-based QC). The phenotypes were cleaned and normalized using the same strategy described above. The first 20 PCs were included as covariates in the fastGWA analysis to control for population stratification. To get a set of independent SNPs associated with a trait, we did a LD-based clumping analysis in PLINK^37^ (threshold *P*-value = 5 × 10^−8^, window size = 1Mb and LD *r*^2^ threshold = 0.01). After the clumping analysis, there were 538 and 57 near-independent SNPs associated with height and BMI respectively, which we call sentinel SNPs. To avoid potential bias in 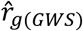 due to remaining LD between the sentinel SNPs, we performed an additional round of the clumping analysis with a much larger window size (i.e., 10Mb) and obtained 531 and 56 sentinel SNPs for height and BMI respectively. The sampling variance of 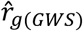 was computed by a Jackknife resampling process^31^.

## URLs

GCTA: http://cnsgenomics.com/software/gcta

PLINK: https://www.cog-genomics.org/plink2

GWAS summary data for height and BMI in GIANT: https://www.broadinstitute.org/collaboration/giant/index.php/GIANT_consortium_data_files

GWAS summary data for BMI in Biobank Japan in NBDC Human Database: https://humandbs.biosciencedbc.jp/en/

UKB consortium: http://www.ukbiobank.ac.uk/

## Data availability

See URLs and Acknowledgements for GWAS summary data and individual data respectively.

## Acknowledgments

This research was supported by the Australian National Health and Medical Research Council (1078037 and 1113400), the Australian Research Council (FT180100186 and FL180100072), and the Sylvia & Charles Viertel Charitable Foundation (Senior Medical Research Fellowship). This study uses AFR data from the database of Genotypes and Phenotypes (dbGaP) (accession numbers: phs000386 for AFR-WHI; AFR-CARe including phs000557.v4.p1, phs000286.v5.p1, phs000613.v1.p2, phs000284.v2.p1, phs000283.v7.p3 for ARIC, JHS, CARDIA, CFS and MESA) and EUR data from the UKB consortium.

## Author Contributions

JY and PMV conceived the study. JY and JG designed the experiment. JG performed statistical analyses under the assistance and guidance from AB, YW, LJ, LY, MEG, PMV and JY. AB, YW, LJ and LY contributed to data preparation. PMV and JY contributed resources and funding. JG and JY wrote the manuscript with the participation of all authors. All authors reviewed and approved the final manuscript.

