## Supplementary File for "Quantifying genetic heterogeneity between continental populations for human height and body mass index"

Guo *et al.*

**Supplementary Figures 1-4**
**Supplementary Table 1**

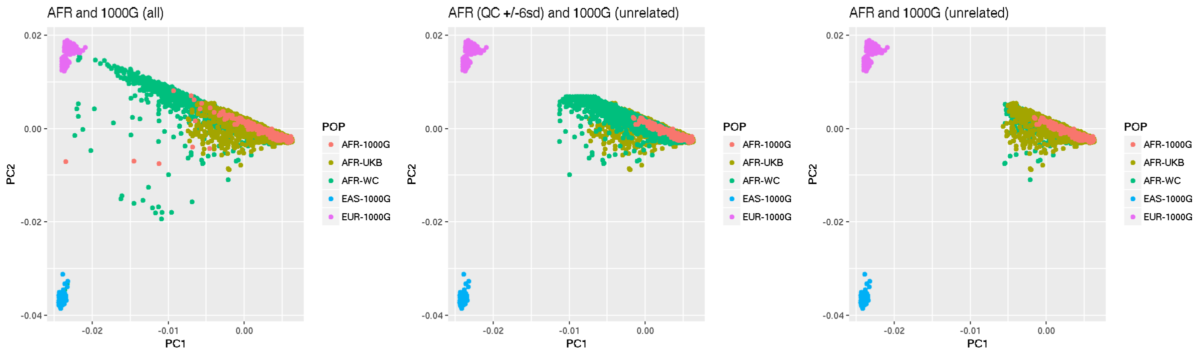


**Supplementary Figure 1** Quality control of the AFR (from UKB and WC; Methods) samples using the principal component analysis based on the common HapMap3 SNPs (MAF > 0.01) in the AFR (from UKB and WC) and the 1000 Genomes Project (1000G) combined samples. The left panel shows all the samples in the 1000G and AFR (from UKB and WC) including relatives, the middle one shows the unrelated individuals (estimated pairwise relatedness < 0.05) for each population in the 1000G and the AFR (from UKB and WC) samples removing the individuals who were more than 6 standard deviations (s.d.) away from the mean of the AFR-1000G and the right panel presents the unrelated individuals in the 1000G and AFR (from UKB and WC). The AFR (from UKB and WC) samples in the middle and right panel were used in fastGWA and GREML analyses respectively (Methods). WC indicates WHI and CARe cohorts (Methods).

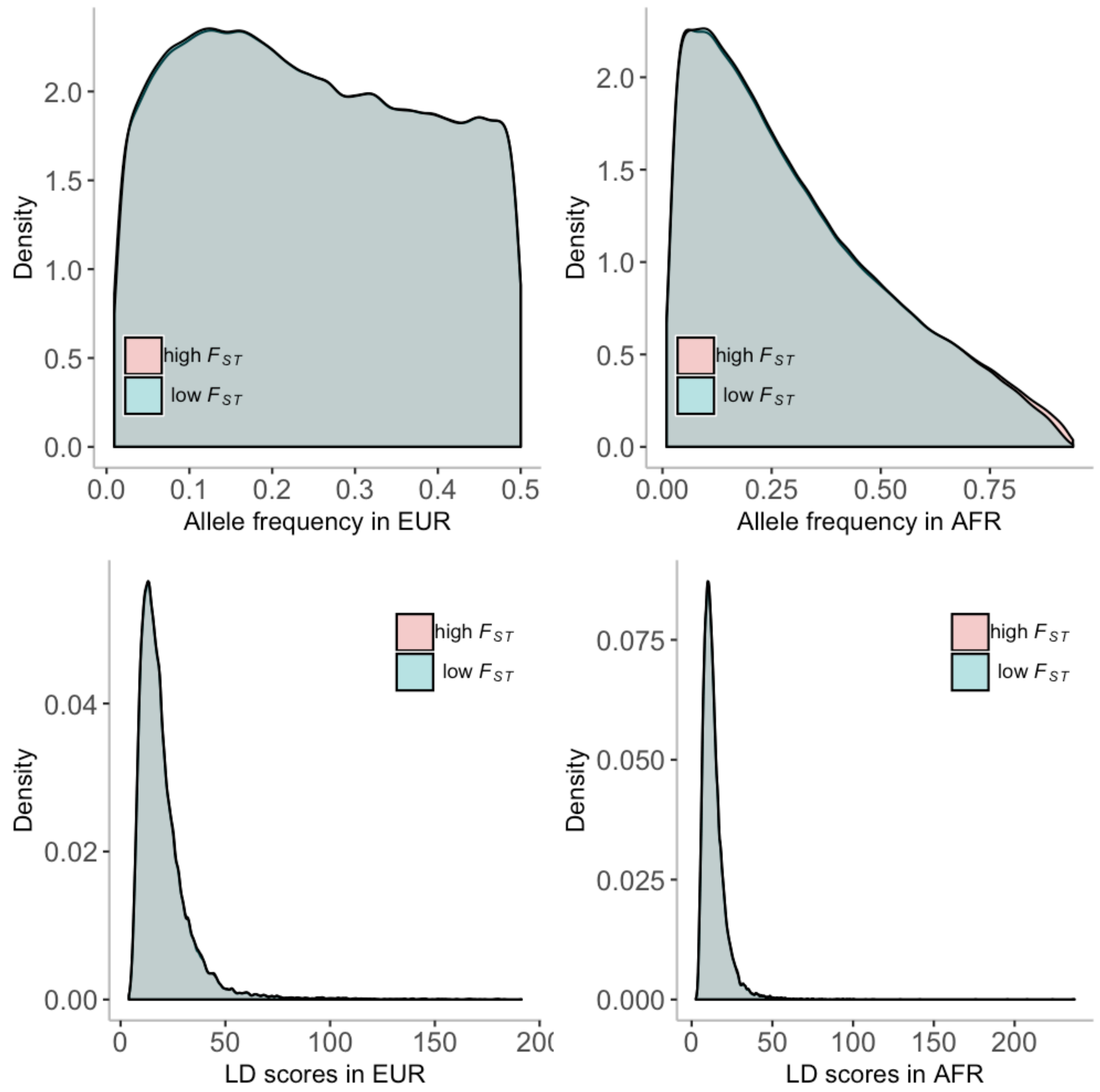


**Supplementary Figure 2** Comparison of the distributions of allele frequencies of the minor alleles in EUR and the same alleles in AFR (upper row), and LD scores in EUR and AFR (lower row) between the SNP sets stratified by $F_{ST}$ values. The two distributions in each panel are almost exactly on top of each other. The LD scores shown here are segment-based LD scores by averaging the per-SNP LD scores across SNPs in a 200kb window with 100kb overlap between two adjacent segments (Methods).

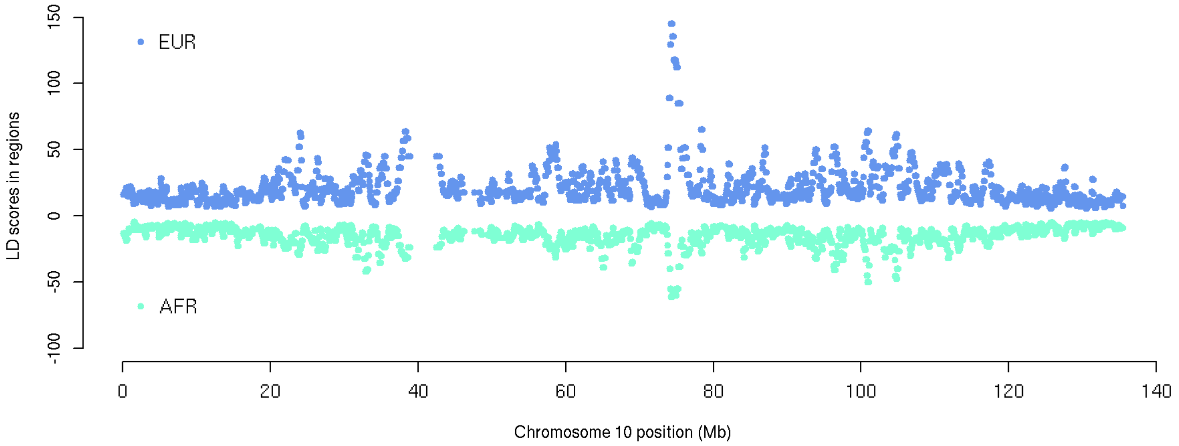


**Supplementary Figure 3** An example plot of the LD scores of the HapMap3 SNPs on chromosome 10 estimated in the unrelated EUR and AFR individuals. Each dot indicates the value of a segment-based LD score by averaging the per-SNP LD scores across SNPs in a 200kb window with 100kb overlap between two adjacent segments (Methods).

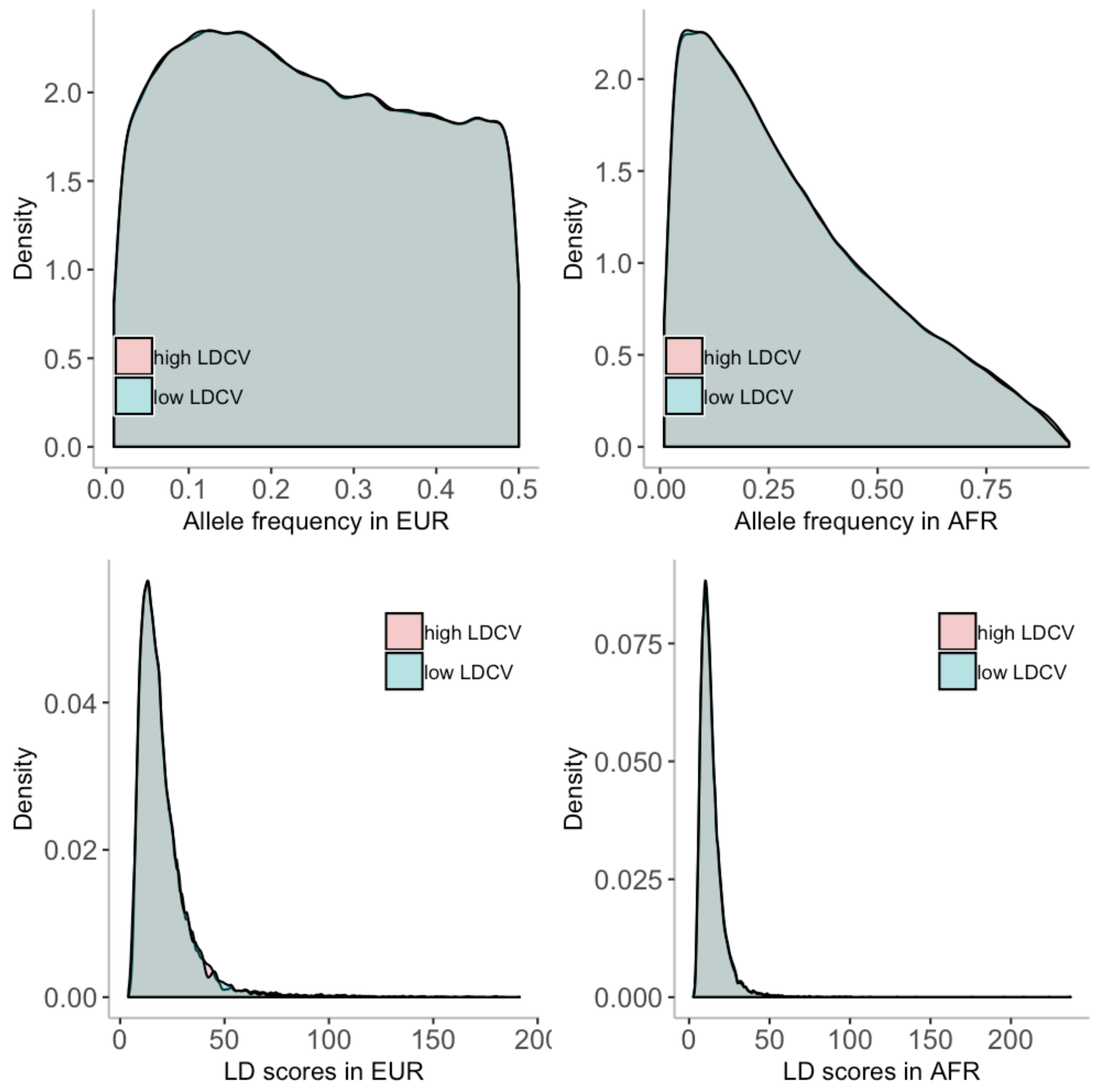


**Supplementary Figure 4** Comparison of the distributions of allele frequencies at the minor alleles in EUR and the same alleles in AFR (upper row), and LD scores in EUR and AFR (lower row) between the SNP sets stratified by LDCV values. The two distributions in each panel are almost exactly on top of each other. The LD scores shown here are segment-based LD scores by averaging the per-SNP LD scores across SNPs in a 200kb window with 100kb overlap between two adjacent segments (Methods).

**Supplementary Table 1** The estimated $\hat{r}_g$ between EUR and AFR using HapMap3 SNPs based on the GRMs-average for height and BMI.

| | $\hat{h}^2_{\mathrm{EUR}}$ **(s.e.)** | $\hat{h}^2_{\mathrm{AFR}}$ **(s.e.)** | $\hat{r}_g$ **(s.e.)** |
|---|---|---|---|
| **Height** | 0.51 (0.0076) | 0.39 (0.024) | 0.78 (0.036) |
| **BMI** | 0.26 (0.0081) | 0.21 (0.024) | 0.72 (0.067) |
